# Bodily perception links memory and self: a case study of an amnesic patient

**DOI:** 10.1101/2025.03.05.640721

**Authors:** Nathalie H. Meyer, Mariana Babo-Rebelo, Jevita Potheegadoo, Lea Duong Phan Thanh, Juliette Boscheron, Bruno Herbelin, Loup Vuarnesson, Sara Stampacchia, Iris M. Toye, Fabienne Esposito, Marilia Morais Lacerda, Arthur Trivier, Elena Beanato, Vincent Alvarez, Michela Bassolino, Olaf Blanke

**Author notes:** Corresponding author: Prof. Olaf Blanke. Equal contributions (Nathalie H. Meyer, Mariana Babo-Rebelo).

## Abstract

Episodic autobiographical memory (EAM) is a building block of self-consciousness, involving recollection and subjective re-experiencing of personal past experiences. Any life episode is originally encoded by a subject within a body. This raises the possibility that memory encoding is shaped by bodily self-consciousness (BSC), a basic form of self-consciousness arising from the multisensory and sensorimotor perceptual signals from the body. Recent studies in healthy subjects showed that embodied encoding improves EAM, with the involvement of the hippocampus. However, there are only few imaging studies to date, hippocampal data are not consistent, and the role of hippocampal damage is not understood. We investigated how different BSC states during encoding, modulate later EAM retrieval, in a patient with severe amnesia caused by rare bilateral hippocampal damage. We performed three separate behavioral experiments using immersive virtual reality. The patient showed consistently more difficulties recollecting information encoded in embodied vs. disembodied states, particularly when asked to recall her perspective experienced at encoding. These results contrasted with the usual beneficial effect of BSC on EAM, and significantly differed from controls. These data provide consistent evidence that BSC impacts encoding and later reliving, and shows that the hippocampus is not just a critical structure for EAM, but also for effects of embodiment on memory. Additional fMRI data extend these findings by revealing that hippocampal-parietal connectivity mediates BSC-EAM coupling. Our findings plead for an important role of BSC in EAM, mediated by the hippocampus and its connectivity, leading to embodied memories that are experienced as belonging to the self.

**Highlights:**

- We studied a patient with memory deficits caused by bilateral hippocampal atrophy
- Multisensory stimulation modulated embodiment during episode encoding
- Across three studies the patient showed stronger memory deficits for embodied episodes
- Multisensory bodily perception impacts bodily self-consciousness and episodic memory
- Our findings link the bodily self to episodic memory and the hippocampus

## Introduction

Episodic autobiographical memory (EAM) refers to memories of contextually rich events from one’s own life (Tulving, 1985) that constitute one’s personal history. It is a building block of self-consciousness, contributing to the maintenance of self-identity across time (Markowitsch and Staniloiu, 2011; Prebble et al., 2013; Tulving, 1985a; Wang, 2011). All life events are composed by a specific content (what), a spatial (where) and a temporal (when) context, that are commonly investigated in EAM research (i.e., Holland and Smulders 2011). Importantly, in addition to these components, all life events are also intrinsically linked to the *subject* or observer who experiences the event, from within the body. Accordingly, the body plays a crucial role in encoding an event. First, it is the support of various multisensory and sensorimotor signals perceived during the encoded event. Second, the integration of these bodily signals gives rise to a form of self-consciousness known as bodily self-consciousness (BSC), (Blanke et al., 2015; Blanke and Metzinger, 2009; De Vignemont, 2011; Ehrsson, 2007; but see next paragraph for detailed introduction). BSC may be particularly relevant for the capacity to develop autonoetic consciousness (ANC) at recall, i.e. to *subjectively re-experience* the sensory details and mental states associated with an event (Tulving, 1985; M. A. Wheeler et al., 1997), a concept tightly linked to self-consciousness. However, despite a growing interest for the understanding of the role of the self, particularly of BSC, in shaping EAM and ANC (Bergouignan et al., 2014; Bréchet et al., 2019; Iriye et al., 2024; Iriye and Ehrsson, 2022; Meyer & Gauthier et al., 2024; Meyer et al., 2024), its behavioral and neural mechanisms and implications are not yet understood. As previous studies mainly focused on the BSC-EAM association in healthy participants, the investigation of clinical cases may bring new and unique insights into the fundamental role of BSC in memory encoding and its neural underpinnings. We investigated the rare case of a patient suffering from severe retrograde amnesia affecting EAM, caused by bilateral hippocampal damage, allowing us to study the impact of different levels of BSC during encoding on subjective experience at recall, and the role of the hippocampus, a key memory structure, in this process.

During the last two decades, the body has been shown to play a key role for the self, in particular with the concept of BSC, a bodily form of self-consciousness arising from the multisensory and sensorimotor integration of bodily signals (Blanke, 2012; Blanke et al., 2015; Blanke and Metzinger, 2009; Ehrsson, 2007; Hyeong-dong Park, Catherine Tallon-Baudry, 2014; Tsakiris et al., 2007). BSC is considered to include four building blocks - a sense of ownership (the feeling that the body belongs to oneself), of agency (the feeling that one is in control of his/her own body), self-location (the feeling that the "self" is located within the body boundaries), and first-person perspective (1PP, the feeling of experiencing the world from of one’s body, in a naturalistic perspective). These four elements are profoundly intertwined and provide a unitary experience of BSC and can be disrupted in neurological patients with out-of-body experiences, associated with disembodiment and the experience of perceiving the world from a third person perspective (3PP) (Blanke and Arzy, 2005; Blanke and Castillo, 2007; Blanke and Mohr, 2005; Jeannerod, 2009; Maeda et al., 2013). BSC can also be altered experimentally in healthy participants, by disrupting perception of the body by exposing participants to conflicting multisensory stimulation using VR (i.e., Aspell et al., 2009; Noel et al., 2015). Some studies using such approaches have provided initial evidence for an implication of BSC in the encoding of episodic memories. For example, encoding an event from a 1PP gives rise to stronger episodic memories than encoding an event from the third-person perspective (3PP; Bergouignan et al., 2014; St. Jacques, 2019; St. Jacques, 2024), and encoding under the illusion of being out of one’s own body causes more 3PP at recall (Bergouignan, 2021; Iriye and St. Jacques, 2021). Consistently with these findings, our group showed a decrease in EAM accuracy when an event was encoded without a body view compared to an event encoded from a 1PP (Bréchet et al., 2019) or when encoded from 3PP versus 1PP (Meyer & Gauthier et al., 2024). These behavioral results provide initial insight into the impact of BSC on EAM.

The hippocampus, and particularly the hippocampal-neocortical axis, is a key candidate for supporting BSC-EAM interactions. This region plays a critical role in EAM and current models consider that this system acts as a mediator during the recall of past events, reactivating cortical areas recruited at encoding (Moscovitch et al., 2016; Nadel and Moscovitch, 2001; Sekeres et al., 2018, 2017). Numerous clinical cases have highlighted the crucial role of the hippocampus (and the prefrontal cortex) in EAM, consistently reporting decreased memory abilities (spatial, episodic and subjective) in patients with lesions in these regions (Gomez et al., 2012; Illman et al., 2011; Klein and Nichols, 2012; Squire, 2009). Critically, recent evidence in healthy participants shows that BSC manipulations at encoding interfere with hippocampal activity at recall. Thus, altered BSC states at encoding lead to reduced functional connectivity between the right hippocampus and the right parahippocampus following encoding (Gauthier et al., 2020), and the intensity of activation of the left posterior hippocampus at recall correlates with a decrease in visual vividness ratings (Bergouignan et al., 2014). More recently, our group has shown that hippocampal reinstatement was also dependent on BSC state at encoding (Meyer & Gauthier et al., 2024), suggesting that the bodily context and its related subjective experience during the encoding of an event modulates the memory trace and contributes to the modulation of hippocampal activity between encoding and recall.

The deficit of the patient we describe in this study was specific to EAM and did not affect semantic autobiographical memory, i.e. memories of general self-knowledge (Tulving, 1985), similarly to other clinical cases (De Renzi et al., 1997; Illman et al., 2011; KAPUR et al., 1992; Levine et al., 2009, 1998; Mayes et al., 2003; Piolino et al., 2005). The patient also showed normal mental states of BSC as tested experimentally and normal overall cognitive abilities. In a previous study (Meyer & Gauthier et al., 2024), we reported preliminary data that this patient had *better* episodic memory for scenes that were encoded in an *altered-disembodied* BSC state, contrarily to the usual beneficial effect of preserved and embodied BSC (Bergouignan et al., 2014; Bréchet et al., 2020, 2019; Gauthier et al., 2020; Iriye et al., 2024; Iriye and Ehrsson, 2022; Meyer & Gauthier et al., 2024). Here we designed three new experiments using VR and further investigate in this patient how BSC affects EAM, in terms of memory accuracy and autonoetic consciousness.

We used immersive VR to perform three experiments over a period of 7-24 months after the patient’s initial hospitalization. We induced different types of conflicting multisensory stimulation (mismatch) to manipulate BSC: visuomotor and perspectival mismatch (Experiment 1), visuotactile mismatch (Experiment 2), and perspectival mismatch (Experiment 3). These manipulations induced embodied and disembodied states, i.e. states where the patient embodied or not the body she would see. While Experiment 1 involved passive and static scenes, Experiment 2 involved life-like scenarios and Experiment 3 required the patient to actively participate in a rich, life-like, social, virtual scenario. We also tested a group of age- and gender-matched control participants in Experiment 1. Given that the patient’s lesions spared BSC-related brain regions, we hypothesized and tested that BSC and the responses to BSC manipulations were comparable to healthy participants. One week after each experiment, we measured different aspects of EAM and autonoetic consciousness, for scenes encoded under Embodied versus Disembodied conditions. In particular, we tested the patient’s ability to retrieve the perspective over a particular scene she experienced at encoding, measuring her ability to re-experience the spatial arrangement around and relative to herself. The one-week interval we applied here was longer relative to previous studies, thereby tackling long-term memory and approaching conditions of autobiographical and life-like memories. We also performed resting-state connectivity analyses, in the patient and a group of controls, searching for hippocampal connectivity patterns that would relate to BSC-EAM coupling.

## Methods

### The patient

The patient was a French speaking right-handed woman in her sixties. She completed the mandatory school years, after which she followed a traineeship. She was working full-time, until her hospitalization. She had no known memory or neuropsychological disorder prior to the brain infection reported in this study.

The patient provided written informed consent, following the local ethical committee recommendations (Cantonal Ethical Committee of Vaud and Valais: 2016-02541) and the declaration of Helsinki (2013).

### Case history

As reported previously (Meyer & Gauthier et al., 2024), the patient experienced epileptic seizures linked to the onset of meningoencephalitis, which developed as a complication of right sphenoidal sinusitis with associated sphenoidal bone loss and meningeal contact. An initial MRI revealed significant lesions in the bilateral medial temporal lobes, affecting both hippocampi, parahippocampi, and amygdalae. Additionally, three smaller lesions (each < 1 cm), were identified in the right middle frontal gyrus, left inferior parietal gyrus, and right lingual gyrus. Follow-up imaging demonstrated gradual improvement, with persistent lesions in the medial temporal lobe and parietal cortex, while the lesions in the lingual gyrus and frontal cortex had resolved by January 2022. The volume of her hippocampi measured eight months after hospitalization was drastically reduced, as reported in (Meyer & Gauthier et al., 2024).

### Neuropsychological assessment and description

Clinically, the patient was diagnosed with moderate to severe retrograde amnesia and moderate anterograde amnesia following a fungal brain infection. Neuropsychological evaluations conducted at the hospital revealed a pronounced deficit in her autobiographical memory, particularly in its episodic content. After her hospitalization, the patient reported initially failing to recognize her apartment, where she had been living for nearly a year. She also experienced topographical orientation difficulties, struggling to navigate familiar environments. Regarding her autobiographical episodic memory (EM), she noted partial improvement in recalling work-related information but continued to have significant gaps in remembering major family events, such as births, marriages, vacations, and deaths. While she recognized familiar faces, she was unable to recall names or the nature of her relationships with those individuals. She was also aware of frequent memory lapses in recent conversations, as close friends pointed out her tendency to repeatedly ask the same questions.

To address these challenges, she began outpatient neuropsychological EM rehabilitation sessions (once weekly from April to June 2022). During these sessions, she underwent several neuropsychological assessments (see Meyer & Gauthier et al., 2024) and had a Montreal Cognitive Assessment (MOCA; Nasreddine et al., 2005) score of 25. However, the primary goal of those sessions was to reconstruct her autobiographical memory by creating a "life axis" using personal photos and anecdotes shared by the patient, and her family and friends.

Despite this structured rehabilitation, her EM deficits persisted. During follow-up sessions and interviews, she struggled to recall autobiographical events and could only provide a limited number of personal semantic memories, which were also impaired. After three months of rehabilitation, she showed slight progress in organizing her life axis and identifying certain life elements. However, she remained unable to vividly evoke or relive any significant life events, even when prompted by others. To support her memory reconstruction, she relied on a "life booklet", collaboratively created with her family and friends, which documented key events in her life.

Trained neuropsychologists from our lab retested the patient at different posterior dates, closer to our experiments, to check for potential improvements or deteriorations of her cognitive abilities (See Supplementary Table 1 for a summary). Briefly, eight months after hospitalization (i.e., one month after Experiment 1), the patient showed preserved executive functions as tested by the Frontal assessment Battery (FAB, Dubois et al., 2000), and the Trail Making Test (TMT, Reitan and Wolfson, 1995). Her episodic memory was along the normal range when tested with the Rey Auditory Verbal Learning Test (RAVLT, Rey, 1964). However, she showed a deficit of EAM, as she recalled few personal events, with little details when assessed with the Autobiographical fluency test (Dritschel et al., 1992). She was not able to mentally relive them, and could only experience a feeling of familiarity when the Remember/Know/Guess paradigm was applied to each (**Table 1**). It is noteworthy that the patient could only guess that she lived certain events after her 30’s, because she saw photographs or was told about them. She was not able to consciously recollect any information about these events.

**Table 1.** Summary results of the autobiographical fluency test. The patient was below the norm for personal episodic content and within the norm for personal semantic information. SD: standard deviation; R: Remember; K: Know; G: Guess; y.o: years old.

| Autobiographical fluency test | Life periods |  |  |  |  |
| --- | --- | --- | --- | --- | --- |
|  | 0-17 y.o. | 18-30 y.o. | >30 y.o. | Last 5 years | Last 12 months |
| Personal semantic information | 23 | 11 | 12 | 10 | 4 |
| Mean (SD) of subjects of the same age | 14.92 (5.28) | 12.83 (9.67) | 13.50 (5.38) | 10.83 (3.78) | 5.58 (4.64) |
| Personal episodic | 1* | 5* | 6 | 0*** | 2 |
| Mean (SD) of subjects of the same age | 7.00 (3.13) | 7.25 (1.76) | 6.00 (1.95) | 4.92 (2.02) | 4.50 (2.07) |
| R/K/G paradigm | K | K | G |  | R |

Eleven months after hospitalization (i.e., one month after Experiment 2), she showed preserved verbal memory (phonemic & semantic verbal fluency test) and episodic memory score within normal range according to the RAVLT, but a deficit in executive functions and visuo-perspective abilities (tested with the Rey Complex Figure test (Rey, 1941), as well as deficit in working memory (tested with the Letter Digit Sequence test). Finally, twenty-four months after hospitalization (i.e., same day as Experiment 3), verbal memory and fluency were normal, she had no signs of apathy, but her frontal efficiency was slightly below normal scores (Supplementary Table 1).

Altogether, these neuropsychological assessments indicate that the patient had remaining EAM deficit at the time of our experiments. The deficit seemed specific to EAM, with preserved semantic memory abilities.

### Experimental procedure

To investigate how hippocampal damage and the related amnesia of the patient may have altered the association of the bodily self with episodic memory and autonoetic consciousness, we performed three experiments in which we manipulated BSC during the encoding of different scenes. **Figure 2** depicts the timeline of our study, from disease onset. In Experiment 1, we manipulated BSC during encoding by exposing the patient to a visuomotor and perspectival mismatch, using VR and a motion tracking system projecting into a virtual avatar which reproduced (or not) the patient’s movements (**Figure 3A**). In Experiment 2, we exposed the patient to a passive visuotactile mismatch, by projecting in front of the patient her own image filmed from behind, while her back was stroke synchronously or asynchronously with the projected image (i.e. out of body illusion). In Experiment 3, we used mixed reality to induce a perspectival mismatch, by showing the patient’s body in 1PP or 3PP, the latter consisting in a 90-degree view of her body (**Figure 6A**). One week after the encoding of each task, we tested the patient’s memory and autonoetic consciousness for different elements of the scenes (**Figures 3B, 4B, 6B**). The following section describes the details of the experimental design used for each experiment.

**Figure 1.**
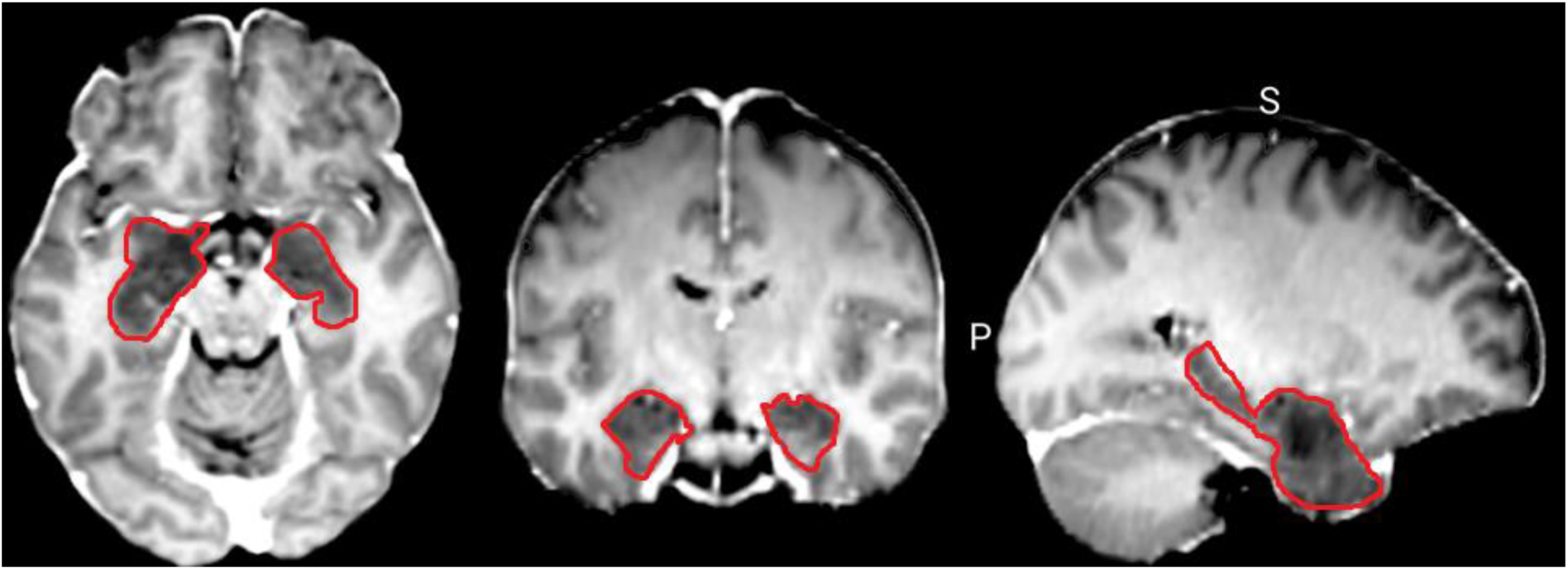
Patient’s anatomical scan. Bilateral inflammation (circled in red) in the amygdalo-hippocampal complex on the day of the diagnosis. S: Superior; P: Posterior; L: Left.

**Figure 2:**
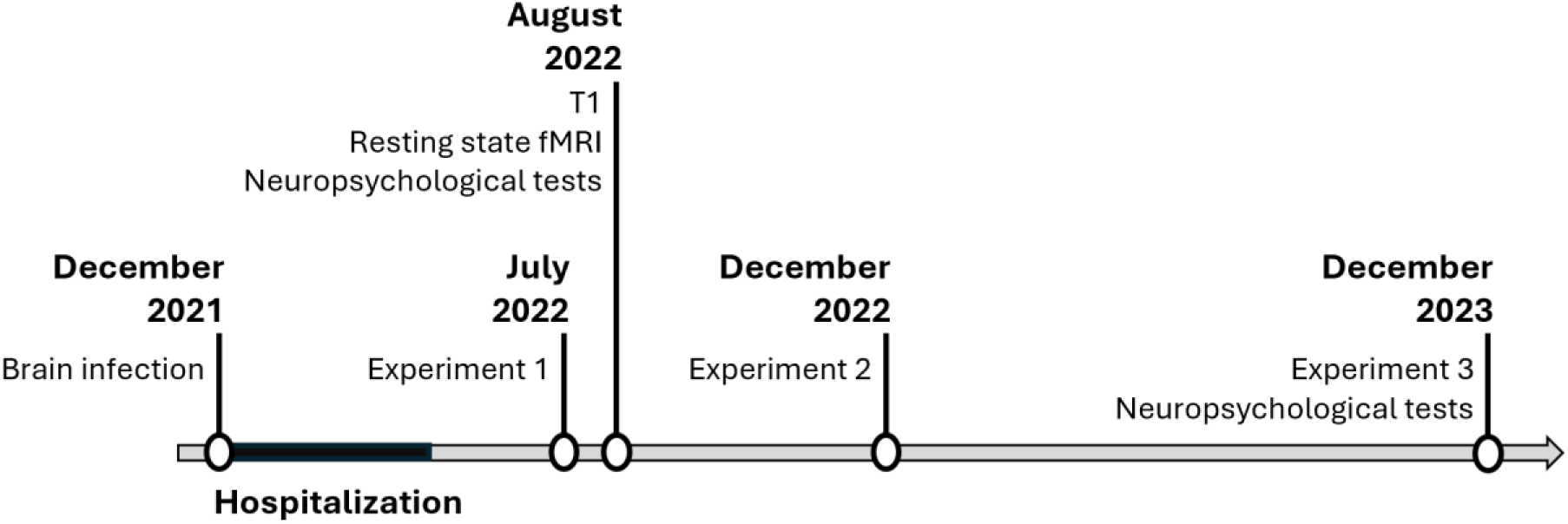
Experiment timeline.

**Figure 3:**
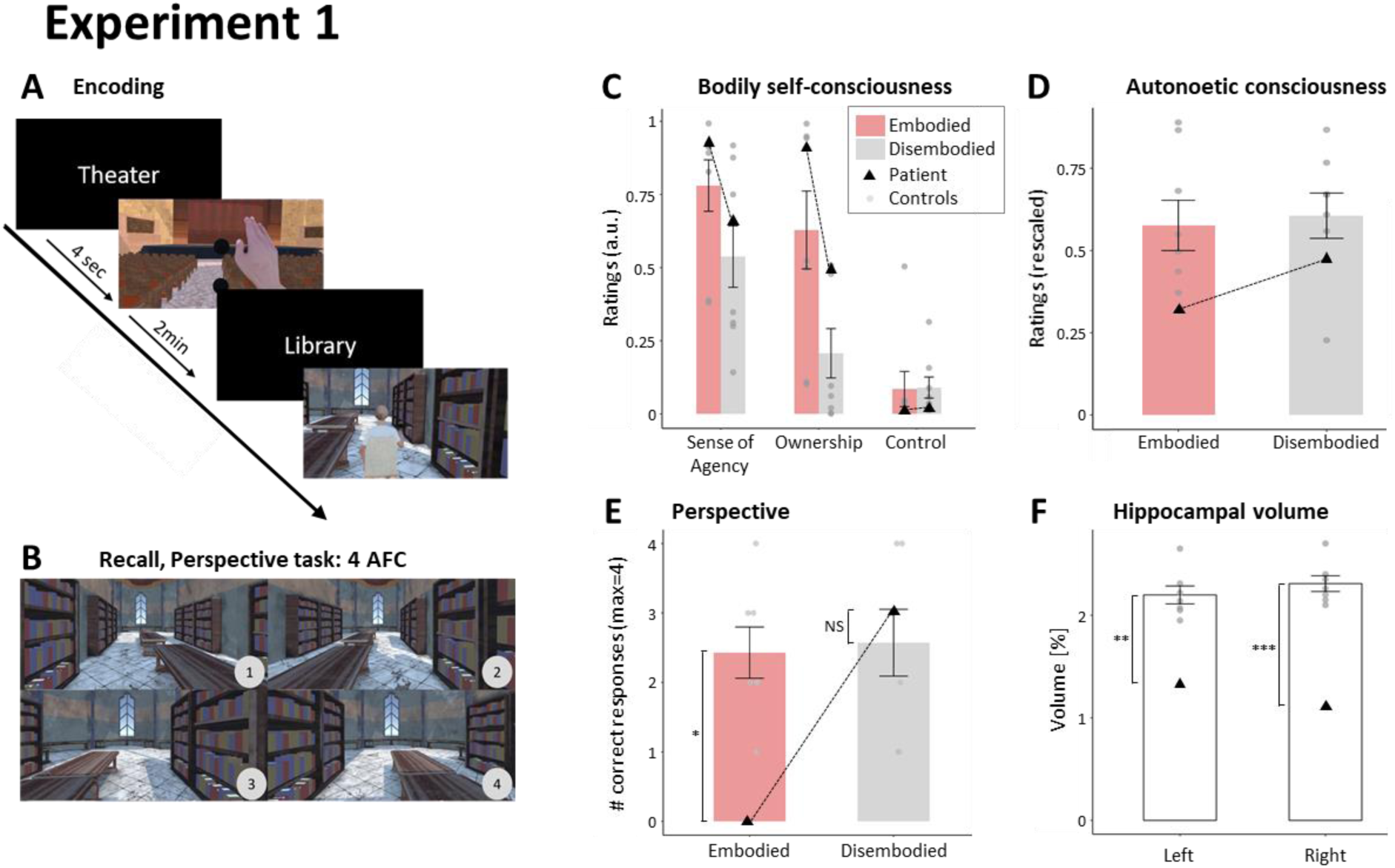
Experiment 1. **A, Encoding.** The patient encoded eight different virtual scenes, where she either viewed an avatar from the first-person perspective moving in synchrony with her own movements (Embodied condition, here “Theater” scene), or an avatar from the third-person perspective whose movements were delayed relative to the patient’s movements (Disembodied condition, here “Library” scene). **B, Recall: 4-alternative forced choice (AFC) task on encoding perspective.** The patient had to choose, among 4 alternatives, the view in which she encoded each scene, here the library scene. **C, Sensitivity to the manipulation of bodily self-consciousness.** The patient (black triangles) had a higher sense of agency and sense of ownership under the Embodied condition, compared to the Disembodied condition, as rated on a continuous scale between 0 and 1. The patient did not respond positively to any of the control questions, accounting for suggestibility. Controls (individual subjects: dots; group average: bars) showed a similar pattern of results. **D, Autonoetic consciousness about the scenes.** The patient remembered fewer scenes and her ratings of autonoetic consciousness (triangles) were lower than controls’ (dots). **E, Perspective task results.** The patient selected the correct perspective for three out of four scenes encoded under the Disembodied condition but did not select any correct perspective for the scenes encoded under the Embodied condition. **F, Hippocampal atrophy.** Both left and right hippocampi of the patient showed reduced volume compared to matched controls. Volume was computed as the percentage of the intracranial volume occupied by the hippocampi. Statistics were computed using a Crawford test, comparing patient against the control group. *: p<0.05; **: p<0.01; ***: p<0.001.

### Experiment 1

#### Encoding

The patient performed Experiment 1 eight months after hospitalization. The patient was successively immersed into eight virtual scenes using a head-mounted display (HMD; Oculus Rift S; refresh rate 80Hz; resolution 1280 x 1440 per eye; 660 ppi), while sitting on a chair and her arm movements being tracked by a motion tracking system (LEAP motion; Leap Motion Controller®; 2 cameras and 3 infrared LEDs) mounted on the HMD. These scenes represented static views of indoor (bedroom, bathroom, kitchen, living room, library, theater) and outdoor (city, stadium) spaces, with a background sound and no human presence. We instructed the patient to move her right arm between two black virtual spheres when they appeared in the scene and to mindwander out loud, including describing the scene and reporting any thoughts or associations that would come to mind (**Figure 3A**). Half of the scenes (bedroom, theater, kitchen and city) were seen from a first-person perspective (1PP), where the position and movements of the virtual arm matched the position and movements of the patient’s arm (Embodied condition: the avatar’s movements and appearance were congruent with the patient’s so the avatar was *embodied*). In the remaining scenes, we induced a visuomotor and perspectival mismatch: a virtual avatar was located 2 virtual meters ahead of the virtual point of view, and its arm movements were delayed (continuous delay varying between 800ms and 1000ms every 5 seconds) compared to the patient’s movements, resulting in a third-person perspective (3PP) view with asynchronous body movements (Disembodied condition: the avatar’s movements and appearance did not match with the patient’s so the avatar was not embodied, it was *disembodied*). This manipulation was shown to decrease the sense of agency and ownership for the virtual body (Meyer et al., 2023). Each scene was encoded for 2 min, and included 3 sequences of 30 seconds of arm movements. Each sequence was followed by 15 seconds of rest during which the patient observed the scene without moving her arm, to avoid tiredness. The order between conditions was randomized.

#### Measure of the strength of BSC manipulation

Right after the encoding session, we tested the patient’s sensitivity to the BSC manipulation by re-testing the same Embodied and Disembodied conditions, in a new outdoor scene (representing a forest). Each condition was shown in 2 blocks of 30s. After each block, the patient rated on a continuous scale her sense of agency ("I felt that I was controlling the virtual body") and her sense of ownership ("I felt that the virtual body was mine"). She also completed two additional questions, controlling for experimental and suggestibility bias ("I felt that I had more than three bodies" and "I felt that the trees were my body").

#### Recall

One week after the encoding session, we tested the patient’s episodic memory for each of the eight virtual scenes in two steps. First, the patient was asked to recall the scene freely, and describe as many elements as she could. When the patient could not directly retrieve the scene, we presented 4 views of the scene on a screen, taken from 4 different perspectives. The avatar was removed from the scene to keep each scene neutral regarding the experimental BSC condition. When this was not sufficient, we replayed the background sound associated with the scene. After this free recall part, we assessed the patient’s autonoetic consciousness for each scene, using an in-house questionnaire (Meyer & Gauthier et al., 2024). This questionnaire included questions from the memory characteristic questionnaire (Johnson et al., 1988) and from the episodic autobiographical memory interview part B (Irish et al., 2011). Second, the patient performed the perspective task (**Figure 3B**), where she indicated which of 4 alternative views of the scene (as displayed on the screen during recall) corresponded to the perspective she experienced during encoding (Perspective Task). For this part, we asked the patient to answer even when she was not able to retrieve any details from the scene.

#### Control group

We performed the same experiment with a group of 7 age-matched women (age, mean±SD: 67±5). Participants were right-handed with no history of neurological or psychiatric disorders and no drug consumption in the 48h hours preceding the experiment. Inclusion criteria additionally required a MOCA score superior or equal to the patient (>= 25), to ensure a comparable level of global cognitive abilities. All participants were compensated for their participation and provided written informed consent following the local ethical committee (Cantonal Ethical Committee Vaud and Valais: 2016-02541) and the declaration of Helsinki (2013). The control group performed the exact same experiment as the patient, with the same stimuli and randomizations, thereby serving as a comparison level for the patient’s results.

### Experiment 2

To replicate and deepen the results of Experiment 1, we tested the patient in a second experiment (Experiment 2), three months after Experiment 1 (i.e. 11 months after hospitalization). Our aim was to ensure that the effect observed in Experiment 1 was not specific to the visuomotor and perspectival mismatch and rather reflected a more general BSC effect, extended to different types of multisensory disruptions. For this we used a visuotactile manipulation of BSC to alter the sense of self-location and body ownership, using the Full-Body Illusion (FBI, Lenggenhager et al., 2007; Noel et al., 2015). Additionally, to extend our understanding on the type of content impacted by the dissociation of EAM and BSC, we also implemented complementary tests to measure different characteristics of EAM that were not tested in Experiment 1.

#### Encoding

We took the patient to 8 different locations on our campus (see **Table 2**), which was a new environment for her. In each location, the patient sat on a chair, wearing an HMD (Oculus Rift DK1, resolution of 640x800, horizontal FOV 90°), and we placed a camera (Logitech C510, resolution 1280x720, 30 fps) 2 meters behind her back. The camera filmed the scene and the patient sitting, and the image was projected on the HMD. During the FBI induction, the experimenter stroked the patient’s back with a wooden stick, as in (Lenggenhager et al., 2007), for one minute. For half of the locations, the stroking was synchronous with the visual stroking that the patient saw on the projection of her own back (Embodied condition: visuotactile stimulations were congruent so the projected body was embodied). In the remaining locations, the stroking and the visual projection were asynchronous (500ms delay, Disembodied condition: visuotactile stimulations were incongruent so the projected body was not embodied). Critically in this setup, the conditions differed in the delay between visual and tactile stimulations, but the body views were identical. The synchronous condition (Embodied) leads to a drift in perceived self-location towards the projected back, revealing that the seen body is more embodied than in the asynchronous condition as shown in (Lenggenhager et al., 2007).

**Table 2.** Experiment 2. Memory accuracy for “What-Where-When” questions and Perspective task. Accuracy (correct=1, incorrect=0) for each of the five questions used to test memory related to the spatiotemporal information and content of each scene and for the perspective task. Q1 = What was the gender of the actor? Q2 = Did the actor succeed in his/her task? Q3 = Did the scene take place on the left or right visual field? Q4 = Did the scene take place on the ground or upper floor? Q5 = Was the scene played in the morning or afternoon?

| Scene | Condition | Q1 | Q2 | Q3 | Q4 | Q5 | Perspective Task |
| --- | --- | --- | --- | --- | --- | --- | --- |
| Printer | Embodied | 1 | 1 | 1 | 0 | 0 | 0 |
| Terrasse | Disembodied | 1 | 0 | 0 | 1 | 1 | 0 |
| Lab room | Embodied | 0 | 1 | 0 | 1 | 1 | 0 |
| Coffee machine | Disembodied | 0 | 0 | 0 | 1 | 1 | 1 |
| Auditorium | Embodied | 1 | 0 | 1 | 1 | 1 | 0 |
| Delivery dock | Disembodied | 1 | 1 | 1 | 1 | 1 | 1 |
| Library | Embodied | 0 | 1 | 1 | 1 | 1 | 1 |
| Reception | Disembodied | 1 | 0 | 1 | 1 | 0 | 1 |

Following FBI induction, an actor (represented by lab members, that the patient never met prior to the experiment) entered the camera’s field of view and played a scene for two minutes. In each scene, the actor tried to use an object placed in the scene (e.g., using a videoprojector), while showing difficulties achieving his/her goal. He/She did not interact with the patient. The scenes differed in terms of the gender of the actor, success of the actor’s action (e.g., successfully making a photocopy vs unsuccessfully trying to make a coffee), spatial location of the action in the field of view (right/left visual field), and global spatial location within the campus (ground / upper floor). Importantly, the location of the patient relative to the global environment varied, to avoid having a point of view that was systematically aligned with the main reference points of the environment (i.e. systematically facing a wall vs being positioned obliquely relative to a wall). The eight scenes were spanned along the whole day, 4 scenes being encoded in the morning and 4 scenes encoded in the afternoon. The visuotactile conditions (Embodied and Disembodied) were counterbalanced between scenes (**Table 2**).

#### Post-test: Measure of the strength of BSC manipulation

To ensure that the FBI manipulation modulated the patient’s self-location, we performed a post-test, at the end of the encoding day, where we induced the FBI eight times, in a neutral black room. The patient stood up and watched on the HMD her back being stroked. The visuotactile stimulation was synchronous in half of the trials, and asynchronous (500ms delay) in the remaining half. The order of the conditions was randomized. After each FBI induction, the patient saw a black screen and the experimenter moved the patient backward with small steps. The patient was then asked to walk back to where she thought she was at the time of the FBI induction. The distance between her location during FBI induction and the reported location was measured, to estimate potential drifts of self-location in the direction of the projected back (Lenggenhager et al., 2007).

#### Recall

One week after the encoding session, we tested the patient’s episodic memory for each scene, in four steps. First, the patient was cued with the name of the scene (for example, “coffee machine”) and was asked to freely recall as many details as possible from that scene. Second, we measured her autonoetic consciousness of the scene with eight questions, related to the strength of vividness, visual details, and global and emotional reliving Supplementary Table 2). Third, we asked forced-choice questions about the content and spatiotemporal context of the scene (Holland and Smulders, 2011): the "What" (Q1: was the actor a male or a female?, Q2: did the actor succeed in his/her task?), the "Where" (Q3: did the event happen on the right or left part of your field of view?, Q4: did the event happen on the ground floor or on another floor?) and the "When" (Q5: did the event happen in the morning or in the afternoon?) dimensions of the scenes. The patient rated her confidence on each response using a 7-point Likert-scale (0: not sure of the answer; 6, totally sure of the answer). Fourth, we showed 4 different views of the scene, taken from 4 different perspectives (as in Experiment 1, Perspective Task). The patient chose the one that corresponded to the perspective in which she originally encoded the scene **(Figure 4A)** and reported how confident she was on her response (7-point Likert scale).

**Figure 4:**
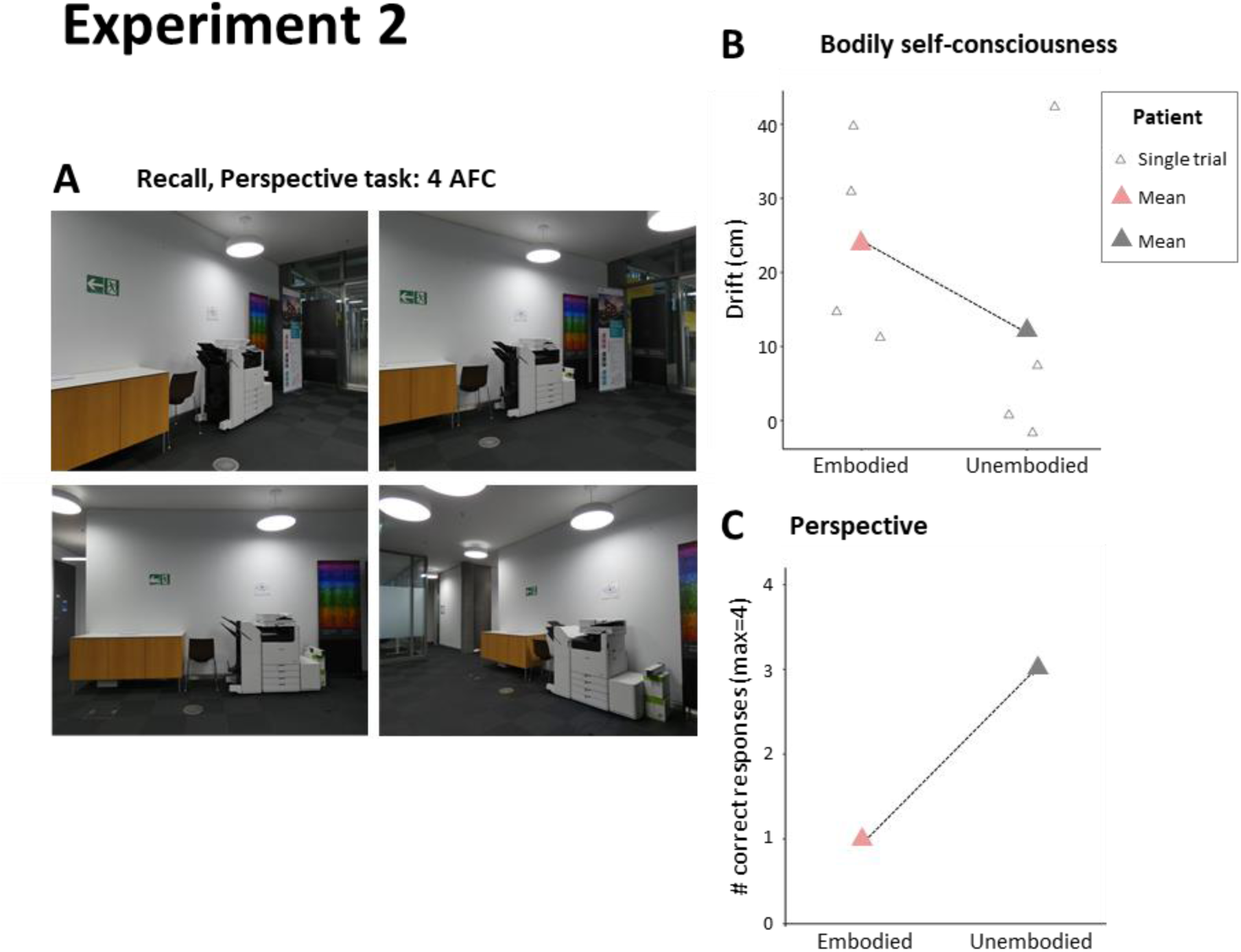
Experiment 2. A, Recall: 4-alternative forced choice task on encoding perspective. The patient had to choose, among 4 alternatives, the view in which she encoded each scene, here the photocopy scene. B, Sensitivity to the manipulation of bodily self-consciousness. The self-location drift following the full-body illusion was larger under the Embodied condition (visuotactile synchrony) than under the Disembodied condition (visuotactile asynchrony). A positive drift indicates a perceived self-location biased towards the observed body’s location, i.e. embodiment of the seen body. White triangles indicate the drift at each trial, and colored triangles indicate the mean across trials for each condition. These measures were obtained during the post-test, at the end of the encoding day (see methods). C, Perspective task results. The patient selected the correct visual perspective for ¾ of the scenes encoded under the Disembodied condition, and only for 1 scene out 4 for the Embodied condition (see also Table 2).

### Experiment 3

In Experiments 1 and 2, the patient passively observed the scenes taking place in front of her. For Experiment 3, we designed a rich immersive experience in mixed reality involving social interactions, body movements and music, contextualized within a story in which the patient was asked to actively interact with the experimenter. We manipulated BSC using a perspectival mismatch. The patient performed Experiment 3 twenty-four months after hospitalization.

#### Encoding

The patient performed the session in a sitting position, while wearing an HMD (HP Reverb G2, resolution of 2160 x 2160 per Eye, FOV = 98.85°) for mixed reality. The mixed reality scenario was built using an in-house software, ExVR (https://github.com/BlankeLab/ExVR). The data of 4 depth cameras (Kinect V2, Microsoft) surrounding and pointing towards the patient and the experimenter was assembled to reconstruct a photorealistic 3D representation of the patient’s and experimenter’s body image which was presented in the VR scene through the HMD in real-time, as in Albert et al. (2019) and Wu et al. (2024).

The patient was first immersed in a virtual replica of the lab room, so that she could familiarize herself with the system and observe her own body. A background voice guided her throughout the experience, which involved a story about the habits and rituals of a community living in nature. The story was divided into four parts, with two BSC conditions (Embodied/Disembodied) and two types of content of the story (about body postures or music), presented in alternation (Embodied/posture, Disembodied/music, Embodied/music, Disembodied/posture). The BSC manipulation consisted in the use of two perspectives, 1PP (Embodied condition: the body seen in mixed reality was embodied) and 3PP (Disembodied condition: the body seen in mixed reality was not embodied), where the patient saw herself from an angle of 90° (**Figure 6A**). Each BSC condition was associated with a distinct guide (blue or orange), a specific view of the virtual environment (forest or mountain) and for the music part, a specific instrument (sleigh bells or maraca). The full story can be read in the supplementary section. The guide sat in front of the patient, so he was seen from the front in the Embodied condition and from a 90° lateral view in the Disembodied condition (**Figure 6A**). The patient was instructed by the background voice to follow the gestures of the guide.

During the body posture parts, the patient had to reproduce the body postures that the guide would perform in front of her. The background voice described and named each posture, and explained its meaning for the community. The guide and the patient performed a series of 4 postures, which they repeated six times.

At the beginning of each music part, the guide gave an instrument to the patient before sitting with his own identical instrument. The background voice explained the importance of the instrument for the community and its making process. The patient was asked to observe it and touch it for one minute. Then, both the guide and the patient accompanied four short music excerpts (30s, the sequence being repeated 2 times) by playing their instruments. After each music excerpt, the patient rated from 1 to 5 how pleasant that music was, by shaking her instrument 1 to 5 times to give her response.

#### Recall

One week after the encoding session, we tested the patient’s memory (**Figure 6B**), in different ways. The first part of the test focused on free recall. The patient was instructed to report everything she could remember about the task (general free recall). Next, the patient completed an autonoetic consciousness questionnaire (questions 16-24 from Experiment 1, as in Experiment 2) for the forest scene (Embodied condition) and another one for the mountain scene (Disembodied condition).

Second, we focused on the body posture parts. We asked the patient to reproduce as many postures as she could, from the encoding session (free recall on body postures). Next, in a body posture recognition task, the patient reported whether each picture we showed her represented a body posture that she performed or not (binary choice; 8 postures performed and 8 new postures, randomized), and her confidence on her response (discrete ratings from 1 to 6, 6 corresponding to a high level of confidence). If she reported recognizing a posture, she had to indicate in which environment she performed it (forest or mountain). We then showed the picture of each of the eight postures she had performed, and the patient chose the corresponding posture name, among four options presented. She was then asked to order chronologically the postures she performed for each condition. Finally, she was asked six 4-alternative forced choice questions, to assess her memory for factual elements of the story on body postures.

Third, we tested the patient’s ability to replace herself in the scene in a perspective task. She was presented with the virtual environment that she encoded, but the view was slightly rotated compared to her encoding viewpoint. She was asked to rotate her viewpoint using a computer mouse until she reached the position in which she experienced the scene during encoding. Hence, contrary to Experiments 1 and 2, here we obtained a continuous measure to quantify her ability to retrieve perspective. The patient performed the task for each scene (forest and mountain, corresponding to two different encoding conditions), twice.

Fourth, we focused on the music parts. We tested her memory for factual elements of the music story and repeated the perspective task. The patient then performed a music recognition task, where she reported (binary answer yes/no) whether each music excerpt had been played at encoding or not (the 8 excerpts from encoding and 16 new music excerpts, randomized).

Finally, the patient completed a 4-alternative-forced choice perspective task (see Experiments 1 and 2), for the forest and mountain environments.

### Behavioral data analysis

#### Autonoetic consciousness

We rescaled each item of the autonoetic consciousness questionnaire (items 8, 9 and 11 were reverse-coded for overall consistency) by dividing each rating by the highest possible rating of each item (to reach a maximal score of 1 for each item). Autonoetic consciousness scores were obtained for each Embodied/Disembodied condition by first averaging all ratings of the questionnaire for each scene, and then averaging across scenes for each condition. The final scores vary between 0 and 1, with higher scores corresponding to a richer recollection.

#### Memory accuracy

We quantified overall memory accuracy in Experiments 2 and 3 by summing the number of correct answers for each scene (cf. Q1-Q5 per scene (only true/false questions: “What Where When” questions for Experiment 2, and recognition and factual information questions for Experiment 3) to obtain a proportion of correct answers.

#### Confidence

In Experiment 2, we summed the confidence ratings given for each question (cf Q1-Q5), for each scene. Confidence ratings were rescaled to obtain ratings between 0 and 1, from low to high confidence.

#### Spatial accuracy of the perspective task (Experiment 3)

To quantify the patient’s accuracy during the continuous perspective task of Experiment 3, we calculated the angle error by computing the mean across trials of the absolute value of the difference between the angle reported and the correct angle under which each scene was observed.

#### Statistical analysis

Statistically supporting the results of a single case is challenging. We thus decided to perform several experiments, considering that a replication of the effect would give strength to the result. In Experiment 1, we compared conditions (Embodied and Disembodied) and hippocampal size between the patient and controls (each hemisphere) using a Crawford test (included in the library “psycho” from R).

### Hippocampal volume and resting state fMRI

We recorded the brain anatomy and resting state functional activity of the patient nine months after disease onset (i.e. one month after Experiment 1). We performed the same recordings on the seven control participants of Experiment 1 (just before their encoding session). During the scan, participants and the patient were asked to fixate a white cross displayed on a grey screen, while trying to minimize blinking. MR images were acquired using a 3T MRI scanner (MAGNETOM PRISMA; Siemens) using a 64-channel head coil.

#### Anatomical MRI and hippocampal volumetry

The MRI session started with a 5 min anatomical imaging, acquired using a T1-weighted MPRAGE sequence (TR = 2300 ms, TE = 2.25 ms, TI = 900 ms, Slice thickness = 1 mm, In-plane resolution = 1 mm × 1 mm, Number of slices = 208, FoV = 256 mm, Flip angle = 8).

Hippocampal volume estimation of the patient and control participants was performed using volBrain (http://volbrain.upv.es) as described in (Meyer, 2023). The reported hippocampal volume corresponds to the percentage of the hippocampal volume with respect to the total intracranial volume (for each participant/patient).

#### Resting-state MRI acquisition and preprocessing

Resting-state fMRI was acquired using a gradient echo planar sequence (TR = 1500ms, TE = 3 ms, Slice thickness = 2mm, number of slices = 69, in-plane resolution = 2 mm × 2 mm, Multiband factor = 3, slice acquisition order = interleaved).

Data was preprocessed using the standard default processing of the conn toolbox version 22a (www.nitrc.org/projects/conn), including slice timing correction, realignment, segmentation, normalization to MNI plane, and smoothing (Nieto-Castanon, 2020).

#### Seed-to-whole brain analysis

To investigate how the hippocampus (atrophied in the patient) could affect the association between BSC and EAM, we used the CONN toolbox to perform two seed-to whole brain analysis, taking as a seed the right and left hippocampus derived from the Harvard-Oxford Cortical atlas (Makris et al., 2006) included in the CONN toolbox.

Functional data were denoised using a standard denoising pipeline (Nieto-Castanon, 2020) followed by bandpass frequency filtering of the BOLD timeseries between 0.008 Hz and 0.09 Hz. Seed-based connectivity maps were estimated characterizing the patterns of functional connectivity with the right and left hippocampus. Functional connectivity strength was represented by Fisher-transformed bivariate correlation coefficients from a weighted general linear model defined separately for each pair of seeds modeling the association between their BOLD signal timeseries. In order to compensate for possible transient magnetization effects at the beginning of each run, individual scans were weighted by a step function convolved with an SPM canonical hemodynamic response function and rectified.

Group-level analyses were performed using a General Linear Model (GLM, Nieto-Castanon, 2020). For each individual voxel a separate GLM was estimated, with first-level connectivity measures at this voxel as dependent variables, and groups or other subject-level identifiers as independent variables. Voxel-level hypotheses were evaluated using multivariate parametric statistics with random-effects across subjects and sample covariance estimation across multiple measurements. Inferences were performed at the level of individual clusters (groups of contiguous voxels). Cluster-level inferences were based on parametric statistics from Gaussian Random Field theory. Results were thresholded using a combination of a cluster-forming p < 0.001 voxel-level threshold, and a familywise corrected p-FDR < 0.05 cluster-size threshold.

Second-level analysis included the 7 control participants from Experiment 1 and the patient. We added the difference in perspective performances between Embodied and Disembodied conditions (Experiment 1) as covariate in the model. Thus, we looked for brain regions whose functional connectivity covaried with the difference in performances in the perspective task, due to the BSC manipulation. We report the coordinate, T-value, cluster size and cluster-size p-value in the Results section, and display the connectivity map on 3D brain render from MRIcroGL using MNI152 template.

To further understand whether the resting-state functional connectivity strength identified with the method above was related to performance in the Embodied or Disembodied conditions specifically, we computed two posthoc Pearson correlations across control participants and the patient, between the connectivity strength results obtained with the above method and the perspective performance for each condition separately.

## Results

All neuropsychological assessments consistently confirmed the patient’s persisting EAM deficit, 24 months after disease onset (for a complete description see “Methods, Neuropsychological assessment and description” and Supplementary Materials). In the following we report the data from the three VR based experiments.

### Experiment 1

#### Validation of BSC manipulation

We first verified that the applied visuomotor and perspectival manipulation induced the expected changes in BSC. The patient’s sensitivity to the manipulation was comparable to that observed in matched healthy controls (**Figure 3C**; Supplementary Table 3) in both conditions. Visuomotor and perspectival mismatch induced a decreased sense of agency and body ownership, compared to the condition with visuomotor and perspectival congruency. Moreover, the ratings on the control questions were low and did not differ between conditions, for both the patient and controls, similar to previous work on healthy participants (Meyer & Gauthier et al., 2024; Meyer et al., 2024). Accordingly, we refer to the two experimental conditions as inducing an Embodied (visuomotor and perspectival congruency) and a Disembodied mental state.

#### Embodiment impairs memory for visual perspective judgements

We next investigated how the manipulation of BSC impacted the patient’s memory. During free recall, while control subjects could easily retrieve all 8 encoded scenes (4 embodied and 4 disembodied), the patient could only partially retrieve 5 out of 8 encoded scenes (of which she retrieved 3 only after receiving additional visual and sound cues); 3 of these 5 scenes had been encoded in the Disembodied condition. The overall autonoetic consciousness (ANC) score of the five scenes was low (M=0.17), and significantly lower compared to healthy controls (**Figure 3D**; Embodied condition: Mean ± Standard Deviation _Control_ = 0.61±0.21, p = 0.042; Disembodied condition: M±SD_Control_ = 0.62±0.20; p = 0.042). Moreover, the patient’s ANC scores were slightly lower for scenes encoded in the Embodied condition (Embodied condition M=0.16, Disembodied condition M=0.18), suggesting even poorer ability to relive scenes that were encoded under in an Embodied versus Disembodied mental state.

In the perspective task and in the Disembodied condition (**Figure 3E**), the patient selected the correct view in 3 out of 4 encoded scenes (chance level per trial: 25%), but gave no correct answer for scenes encoded in the Embodied condition. Her performance in the Embodied condition was significantly lower when compared to the control group (M_controls_±SDcontrols = 2.43±0.98, p = 0.03), whereas performance in the Disembodied condition did not differ from controls (M_controls_±SDcontrols = 2.5±1.27, p = 0.34).

#### A significant reduction in hippocampal volume

The patient had smaller hippocampi than the group of healthy controls, for both hemispheres (**Figure 3F**; left hippocampus: M_controls_ = 2.2%, SD_controls_ = 0.23, p = 0.006; right hippocampus: M_controls_ = 2.3%, SD_controls_ = 0.2, p < 0.001). None of the controls had atrophied hippocampi compared to the database of the toolbox (volbrain) used to perform the analysis. A whole medial temporal lobe analysis did not point to any other differences in regional volume between the patient and controls.

This experiment highlights the patient’s memory impairment relative to healthy controls. The results suggest that the impairment is more pronounced for encoding under an embodied condition, indicating that sensorimotor integration during encoding is processed differently in the patient compared to healthy controls.

### Experiment 2

#### Validation of BSC manipulation

Similarly to what is classically obtained in healthy participants (Lenggenhager et al., 2007; Pfeiffer et al., 2011), the patient showed a positive drift of perceived self-location towards the projected image of her body (measured during the post-test). The drift was larger in the visuo-tactile congruent condition (**Figure 4B**; Mean ± Standard Deviation = 24.25±13cm) than the visuo-tactile mismatch condition (M±SD = 12 ±20cm). This drift in self-location towards the location of the projected body is an indication of the level of embodiment for the projected body. This indicates, as in Experiment 1, that the patient was sensitive to the BSC manipulation and that the congruent visuo-tactile condition enabled stronger embodiment of the seen body (as compared to the visuo-tactile mismatch condition). Accordingly, we refer to the two experimental conditions as inducing an Embodied and Disembodied mental state, respectively.

#### Embodiment impairs memory for visual perspective judgements

One week after encoding, the patient performed the perspective task, where 4 alternative views of each scene were presented (**Figure 4A**) and the patient selected the one that she believed corresponded to the view she experienced at encoding (as in Experiment 1). Consistently with the results obtained in Experiment 1, the patient selected the correct view for three out of four scenes encoded in the Disembodied condition, but gave only one correct answer from the 4 scenes she encoded in the Embodied condition (**Figure 4C**).

#### Embodied condition was associated with decreased feeling of re-experiencing and decreased confidence of remembering

The patient gave higher autonoetic consciousness ratings when recollecting scenes in the Disembodied condition (Supplementary Figure; Supplementary Table 2; M±SD = 0.7±0.2, rescaled scores) compared to scenes encoded under the Embodied condition (M±SD = 0.49±0.1). This difference seemed to be mainly due to the recollection of information related to the global experience (visual details, global level of details, spatial context), rather than to the reliving dimension (global and emotional), which was rated very low for all scenes and did not differ between conditions (Supplementary Table 2).

We tested the patient’s memory for the content and spatiotemporal context of the encoded scenes ("What-Where-When", see Methods), by presenting her five two-alternative forced choice questions for each scene. The patient could remember only part of the information of each scene and this was equal for the Embodied and Disembodied condition (**Table 2**; Supplementary Figure; Supplementary Table 2). However, the patient reported being more confident on her answers about scenes encoded in the Disembodied condition (M±SD = 0.93±0.4), as compared to scenes encoded in the Embodied condition (M±SD = 0.68±0.37; Supplementary Figure; Supplementary Table 3).

Overall, these results confirm and extend the findings of Experiment 1. Once more, the Embodied condition impaired memory for the perspective perceived at encoding. In addition, we show here that despite similar levels of memory accuracy, the feeling of remembering and of re-experiencing (confidence and autonoetic consciousness) was stronger in the Disembodied condition.

### Experiment 3

#### Poor autonoetic consciousness despite richer and more engaging virtual reality scenario

During free recall, the patient remembered having been in a forest and having met different guides. As she did not remember seeing a mountain, we could not measure autonoetic consciousness separately for the forest (Embodied condition) and the mountain (Disembodied condition), therefore we asked her to answer based on her global experience of the virtual reality environment. Even with a much richer virtual reality environment and a more active and engaging story line, her global autonoetic consciousness was low and comparable to autonoetic consciousness in Experiments 1 and 2, as well as consistently lower than controls of Experiment 1 (**Figure 5**). This confirms that her low levels of autonoetic consciousness are a general trait of her amnestic state, and not related to her engagement in the task or to the sensory richness of the experiment.

**Figure 5:**
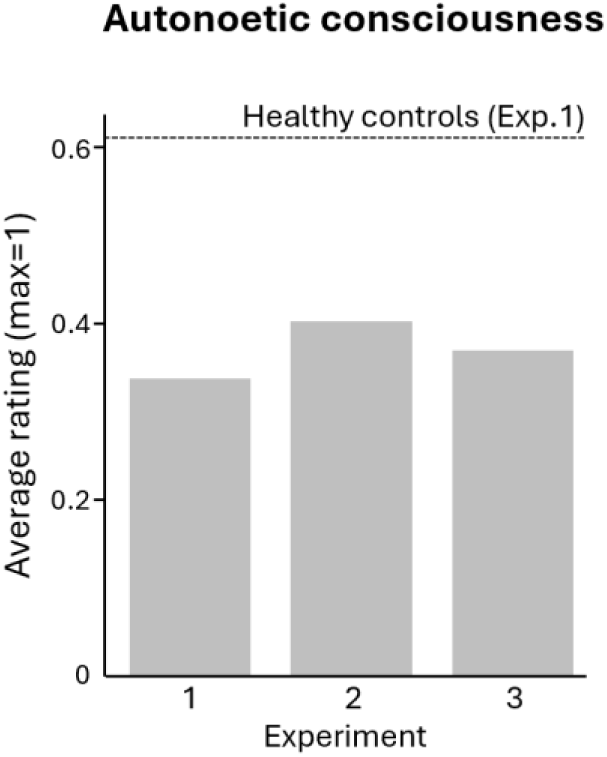
Autonoetic consciousnesss across experiments. The bars correspond the average ratings of the patient, for each of the three experiments. The dashed line represented the average rating obtained for the group of healthy controls (n=7), in Experiment 1.

#### Embodiment impairs memory for perspective and for factual information

Although the patient had difficulties performing our task and reported answering mostly by chance, we found similar results in the perspective task, as compared to Experiments 1 and 2. The patient was more accurate at reporting the viewpoint in the Disembodied condition (mountain) than the Embodied (forest) condition (**Figure 6C**; mean of the angle error (absolute value), across the 2 repetitions of the task: Embodied condition = 33.5°, Disembodied condition = 22°). The patient did not give a single correct response on the 4 alternative-forced-choice version of the Perspective task.

**Figure 6:**
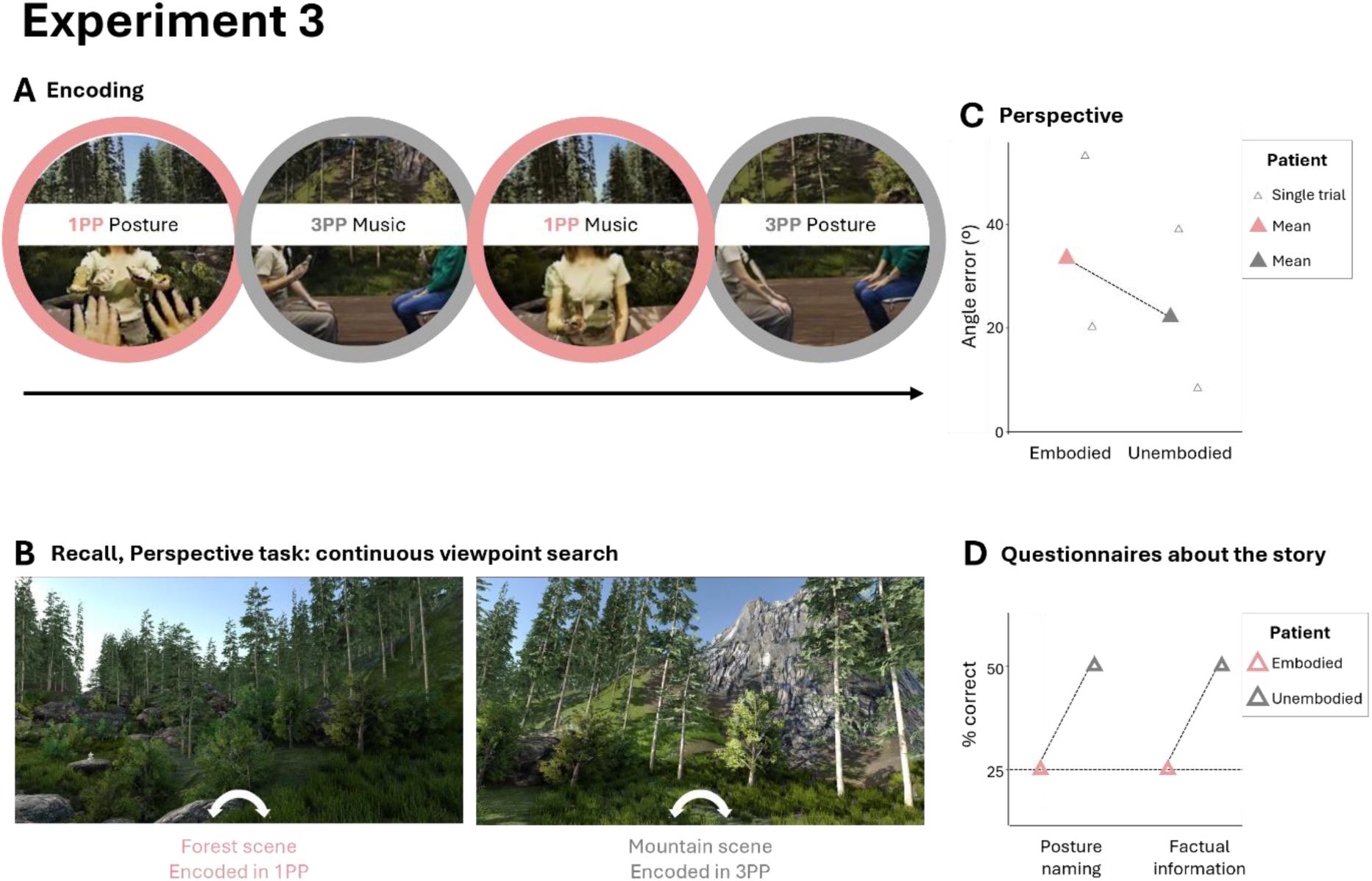
Experiment 3. **A, Encoding.** The patient successively encoded 4 scenes, involving a mixed-reality setup where she would see her body from a first-person perspective (1PP, Embodied condition) or from a third-person perspective (3PP, 90° view shift, Disembodied condition). The scenes involved either playing an instrument with the guide (“Music” scenes) or performing body postures following the guide’s movements (“Posture” scenes). **B, Recall: continuous search of encoding perspective.** In this Perspective task, the patient was shown the virtual environment on a computer screen and was asked to rotate the view until she found her encoding viewpoint. The Embodied condition was associated with a forest environment, while the Disembodied condition was associated with the view of a mountain. **C, Perspective task results.** The patient showed worse precision at reporting the view encoded under the Embodied condition compared to the Disembodied condition. We here represent the absolute value of the angle error in the continuous perspective task (B). White triangles represent individual trials, colored triangles represent the average across trials. **D, Performances on the posture naming and factual information questionnaires.** The patient showed better performances for the Disembodied condition in both tasks (four-alternative forced choice). The dotted line represents chance level.

In the Disembodied condition the patient was again better and, importantly, above chance level, on the questions about the names and factual information of the body postures (**Figure 6D**). The patient did not succeed in reordering the postures performed during either the Embodied or Disembodied blocks. The patient had 75% of hit rate and correct rejection for the postures performed in both Embodied and Disembodied conditions. Additionally, the patient was better at recognizing the music excerpts played during the Embodied than Disembodied condition (Embodied: hit rate = 100%; Disembodied: hit rate = 50%; Correct rejection = 50%).

Together, the results from Experiment 3 further support the consistent finding that the patient has more difficulties retrieving the perceived perspective at encoding when the scenes were encoded in the Embodied condition.

### Resting state fMRI

#### Embodiment modulates functional connectivity between the right hippocampus and regions of the BSC network

In addition to the behavioral data described above, in Experiment 1, we acquired resting state fMRI of the patient and healthy controls, prior to retrieval. We tested whether the magnitude of the effect of the BSC manipulation on the perspective task (i.e. the performance difference between the Embodied and Disembodied conditions) was associated with resting-state functional connectivity between the hippocampus (seeds) and other regions of the brain, for the patient and control group taken together.

We found a negative modulation of the functional connectivity of the right hippocampus with the angular gyrus (**Figure 7A**; MNI coordinates x = -54, y = -58, z = 46, cluster size = 47, pFDR< 0.001), and the frontal pole (MNI coordinates x =12, y = 46, z = 2, cluster size 51, pFDR < 0.001). This finding shows that the stronger the functional connectivity at rest between these regions and the right hippocampus, the smaller the difference of performance between Embodied and Disembodied conditions. We additionally found a positive modulation of the functional connectivity of the right hippocampus with the right superior parietal lobule and middle frontal gyrus (**Figure 7A**; Superior parietal lobule: MNI coordinates x = 24, y = -50, z = 72, cluster size = 34, pFDR < 0.001, see Supplementary Table 4 for the details of each cluster). The same analysis with the left hippocampus did not lead to any significant clusters.

**Figure 7:**
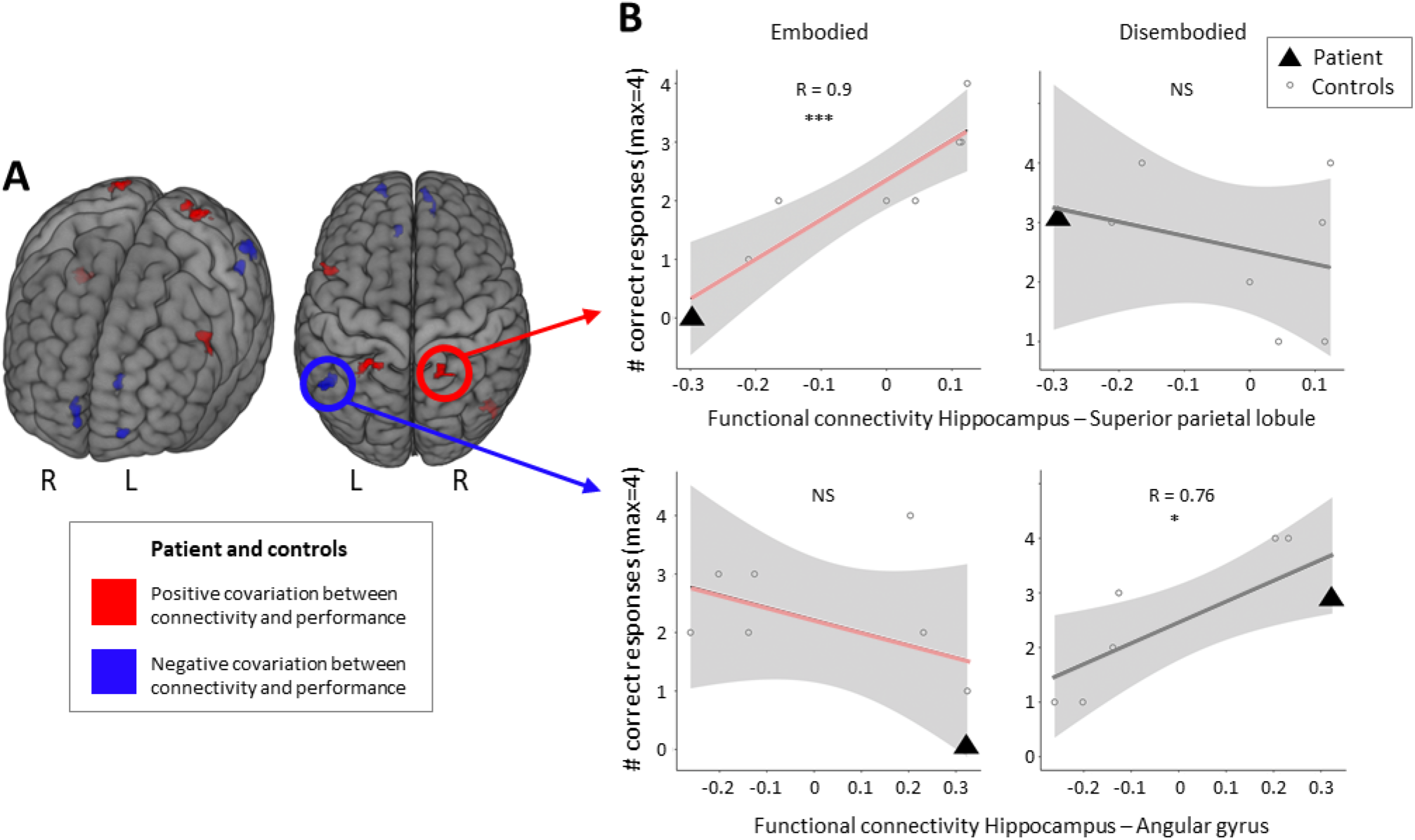
Performance on the Perspective task (Experiment 1) is associated with the functional connectivity between the hippocampus and distinct cortical regions. A, Hippocampal connectivity. Results of the seed (right hippocampus) to whole brain functional connectivity analysis with the difference in perspective performance (Embodied-Disembodied) as covariate, using the resting state fMRI of the control participants (n=7) and the resting state fMRI of the patient (acquired 1 month after Experiment 1). Red clusters depict voxels which connectivity with the hippocampus covaries positively with the difference in perspective scores. Blue clusters depict voxels which connectivity with the hippocampus covaries negatively with the difference in perspective scores. Brain regions were identified using a cluster-forming voxel threshold of p<0.001 and a cluster-size threshold corrected for false discovery rate pFDR<0.05. **B, Connectivity and Embodied vs Disembodied condition**. The two plots on the top represent Pearson correlations between the functional connectivity strength between right hippocampus and right superior parietal lobule, and performance on the Perspective task for Embodied (left) and Disembodied (right) conditions (Experiment 1). Similarly, the two plots on the bottom correspond to right hippocampus – angular gyrus functional connectivity. Black triangles represent the patient, grey dots represent the individual control subjects. *: p<0.05; ***: p<0.001.

We next split performance between Embodied and Disembodied conditions, and tested whether these resting-state functional connectivity results could be better related to any of the two conditions. Better performance in the Embodied condition was associated with higher resting state functional connectivity between the right hippocampus and the right superior parietal lobule (**Table 3)**. The patient showed a lower functional connectivity compared to healthy controls (**Figure 7B**) between these two regions. Conversely, better performance in the Disembodied condition was associated with higher resting state functional connectivity between the right hippocampus and left angular gyrus (**Figure 7B**; **Table 3**). Here, the patient showed a stronger functional connectivity compared to healthy controls. These results suggest that the strength of the functional connectivity at rest between the right hippocampus and different posterior parietal regions is associated with performance in the perspective task, with specific connections being more prominent in the Embodied or Disembodied conditions.

**Table 3:** Correlation between the perspective performance for each condition and the resting state functional connectivity between the right hippocampus and different cortical clusters, across controls and the patient. Each cluster was previously identified as being sensitive to the magnitude of the Embodied-Disembodied effect on the Perspective task (Experiment 1). Significant correlations are indicated in bold (uncorrected for multiple comparisons).

| Node connected with the right hippocampus | Correlations, Embodied condition<br>($r$, $t$, $p$-value) | Correlations, Disembodied condition<br>($r$, $t$, $p$-value) |
| --- | --- | --- |
| Lateral Occipital Cortex right | -0.43, -1.16, 0.29 | **0.79, 3.17, 0.02** |
| Frontal Pole left | 0.54, 1.58, 0.17 | -0.5, -1.44, 0.2 |
| Superior Frontal gyrus right | **-0.74, -2.72, 0.034** | 0.53, 1.55, 0.17 |
| Frontal Pole right | -0.5, -1.42, 0.2 | **0.74, 2.75, 0.03** |
| Anterior Cingulate Gyrus left | -0.17, -0.43, 0.69 | 0.43, 1.19, 0.27 |
| Middle Frontal Gyrus left | 0.63, 2.016, 0.09 | **-0.71, -2.53, 0.04** |
| Angular Gyrus left | -0.67, -2.23, 0.07 | 0.43, 1.18, 0.28 |
| Superior Parietal lobule left | 0.69, 2.39, 0.053 | -0.52, -1.52, 0.18 |
| Superior Parietal Lobule right | **0.9, 5.28, 0.0019** | -0.33, -0.87, 0.41 |

## Discussion

We report the case of a patient with severe episodic autobiographical memory loss, but preserved global cognitive functions, that was caused by bilateral hippocampal atrophy. We manipulated bodily self-consciousness in different ways across three experiments, with the goal of assessing the impact of embodiment on episodic memory encoding in this rare clinical case. We found that the patient showed normal sensitivity to BSC manipulations, comparable to what is usually reported in the literature for healthy participants and comparable to the present control group. Thus, a multisensory match (across the 3 experiments) leads to an *embodied* state, while conditions of multisensory mismatch do not induce such embodiment of the seen body (i.e., *disembodied* state). Concerning episodic memory, we here show a consistent pattern of the patient across three experiments. When encoding was performed under an embodied state the patient showed more difficulties retrieving her viewpoint adopted at encoding, whereas when she encoded similar scenes in a disembodied state, her performance about the encoding viewpoint was better. This effect was accompanied by impairments of autonoetic consciousness for episodes encoded under the embodied state and our imaging finding that the precision of viewpoint retrieval was related to different functional connections between hippocampus and posterior parietal cortex, again depending on the dis-vs. embodied state at encoding. These data provide new evidence for the contribution of bodily self-consciousness to the encoding of life episodes.

### Bodily self at encoding modulates autonoetic consciousness at recall

The three experiments of this study induced various types of multisensory conflicts that modulated different aspects of bodily self-consciousness. In Experiment 1, the sense of ownership and agency were altered following a visuomotor and perspectival mismatch. Experiment 2 applied visuo-tactile stimulation to induce embodiment for the projected body. Experiment 3 created a perspectival mismatch, by which the patient would see herself from the side. The decrease in embodiment experienced by the patient following these manipulations was comparable to what is usually observed in healthy participants (Ehrsson, 2012; Haggard, 2017; Kannape and Blanke, 2013; Lenggenhager et al., 2007) and in Experiment 1 comparable to those of a control group. Therefore conditions involving multisensory stimulations that were congruent with the body, as shown in the visual scene, led to an embodied self within the encoded scene (Embodied conditions). On the contrary, conditions that involved incongruent multisensory stimulations induced a conflict that prevented or decreased an embodied self within the scene during the encoding (Disembodied conditions).

Interestingly, although the patient’s modulation of embodiment as induced by such multisensory bodily stimulations was comparable in strength and sensitivity to healthy participants, the effect of embodiment on memory was opposite to what is usually observed.

The patient showed a detrimental effect of embodiment (or *preserved* bodily self-consciousness) on memory performance (Embodied condition) whereas healthy participants tend to show a positive effect of preserved bodily self-consciousness on memory in previous studies (Bergouignan et al., 2014; Bréchet et al., 2019, 2020; Iriye and Ehrsson, 2022; Meyer & Gauthier et al., 2024). Moreover, the patient’s memory performance was significantly worse in the Embodied condition compared to healthy controls, while her performance in the Disembodied condition was similar to healthy controls, suggesting that her memory was more deficient for scenes in which the self was embodied within the scene compared to scenes where the self was not embodied. We note that the fact that performance did not differ between Embodied and Disembodied conditions for controls should not be taken as a general result, as this group was specifically tested to serve as a baseline level for the performances of the patient. Critically, although each experiment involved only few trials, the patient’s results were replicated in two further separate experiments (Experiments 2 and 3), using different manipulations of embodiment, pointing to a general alteration in how mechanisms of bodily self-consciousness impact episodic memory.

A recent study reported that this patient showed lower performance in a recognition task, when encoding occurred during an embodied mental state ( Meyer & Gauthier et al., 2024). Consistently with this previous result, we here show for the first time that the recollection of the perspective or viewpoint that she adopted at encoding is also affected in conditions with an embodied mental state. The perspective task proved to be the most sensitive to the embodiment manipulation. Performing the perspective task requires to re-visualize the scene, to re-experience the first-person viewpoint of the environment, and in that sense, it revives first-person past experiences (Babo-Rebelo et al., 2022) and engages autonoetic consciousness. Although the patient systematically reported having difficulties performing this task, her performances were better in Experiments 1 and 2 in the Disembodied condition, and the effect was replicated in Experiment 3 with a continuous measure of perspective. Ratings of autonoetic consciousness were slightly lower for the Embodied condition in Experiment 1, and in Experiment 2 the Disembodied condition was associated with a higher feeling of remembering and re-experiencing (higher confidence ratings and higher autonoetic consciousness ratings). The other recall tests in this study were not affected by the present manipulations, likely due to the reduced number of trials.

Retrieving an encoded viewpoint or perspective, the feeling of remembering and the feeling of re-experiencing are all part of the notion of autonoetic consciousness (Gardiner et al., 2001; Tulving, 1985). Our results indicate that the bodily self also has an impact on the autonoetic dimension of memory. Furthermore, the ability to relive an episode and experience autonoetic consciousness has been argued to require retrieving information about this episode, including self-attribution (Tulving, 2002, 1985). Based on the present data we propose that the bodily self is a key component for self-attribution of life events and autonoetic consciousness. We argue that mechanisms of bodily self-consciousness contribute to the encoding of life events as being self-attributed, as belonging to the self. This differs for events encoded under disembodied mental states due to disturbed mechanisms of bodily self-consciousness during encoding that are more detached from the self. The present patient shows a disruption of this coupling between bodily self-consciousness and episodic memory.

### Key role of the bodily self for the self-relatedness of memories: A mechanistic proposal

The fact that the patient shows preserved viewpoint memory for altered bodily self states (Disembodied condition), but impaired viewpoint memory for embodied states (Embodied condition), shows that memory retrieval is modulated by the level of engagement of the bodily self at encoding. Given the specificity of the patient’s deficits to embodied memories when compared to her performance in disembodied condition, the neural mechanisms from retrieval which reflect embodiment during encoding rely on the hippocampal structures that are most affected in the patient.

This is further supported by the results of the functional connectivity analysis (Experiment 1), linking embodiment-related effects on episodic memory encoding to different connectivity patterns between the right hippocampus and posterior parietal cortex, a key area of bodily self-consciousness. More specifically, better perspective performance for memories encoded under *preserved* BSC was associated with a stronger connectivity between the right hippocampus and right superior parietal lobule. The posterior parietal cortex, which includes the superior parietal lobule, is known to play a key role in the multisensory and sensorimotor integration of bodily signals, as evidenced by numerous studies (Iacoboni & Zaidel, 2004; Làdavas & Farnè, 2004; Làdavas & Pavani, 1998; Maravita & Romano, 2018). Specifically, the superior parietal lobule is linked to the sense of body ownership and is a key structure of bodily self-consciousness (BSC) (Ehrsson et al., 2004; Park & Blanke, 2019). Consequently, our data-driven findings underscore the importance of the functional connectivity between memory and BSC-processing regions in accounting memory performance under conditions where BSC is preserved. In contrast, better perspective performance for memories encoded under *disrupted* BSC (disembodied state) was associated with a stronger connectivity between the right hippocampus and the right angular gyrus. Interestingly, the angular gyrus is known for its involvement in episodic memory (Bonnici et al., 2018, 2016; Linden et al., 2017), particularly in its autonoetic component (Bonnici et al 2016; Sestieri et al 2013; for a review Zaman & Russell 2021), but it also has critical role in the first-person perspective (Blanke et al., 2015; Bréchet et al., 2018; Ionta et al., 2011; Martin et al., 2020; van Elk et al., 2017) as well as disembodiment and out-of-body experiences (Blanke et al., 2002, 2004, 2005; Blondiaux et al., 2021). Our results suggest that this connection would be important for memories encoded under a more self-detached state, potentially leading to a distancing of the memory from the self (as has been often reported in memory research (i.e.,Nigro and Neisser, 1983; St. Jacques, 2024, 2022). More work, also in healthy individuals, is needed to understand hippocampal connectivity with both dis/embodiment networks in posterior parietal cortex as well as additional interactions with premotor regions (i.e., Meyer & Gauthier et al., 2024). These experimental results are also consistent with the patient’s episodic autobiographical memory deficit obtained in her clinical routine neuropsychology examinations. These confirmed that the patient’s persistent episodic autobiographical memory deficit which was still present at the time of the present three experiments. The patient was unable to relive past episodes from her life – both before and after the hippocampal damage – although she could still retrieve some semantic information about herself. We also note that the patient’s memory deficits have slightly improved since the onset of her neurological damage, but persist to this day and significantly impact her daily life.

### Limitations

First, it is important to note that this single case report may not be representative of other patients with hippocampal damage who may not show dissociations between BSC and EM. However, our consistent observations highlight the association between the hippocampus and posterior parietal areas in embodied memories. The present paradigms should be tested in other groups of patients with amnesia. Second, this study relies on data collected with a relatively small number of trials per condition. However, we observed consistent results across three different experiments, each engaging different modalities of BSC disruptions, and varying methods of memory assessment. This replication across multiple studies helps mitigate the potential impact of the low trial count per condition. Third, one could argue that Disembodied versus Embodied conditions might have generated stronger changes in attention, thereby creating more favorable conditions for memory encoding. This could potentially explain the better performances of the patient for Disembodied conditions. However, this hypothesis is unlikely, because the experimental manipulations in the three experiments led to highly similar retrieval results although we applied three different multisensory bodily stimulations. While visuomotor and perspectival mismatch can be easily noticeable (Experiment 1), the effect of the full-body illusion in Experiment 2 is much more subtle and not necessarily consciously perceived. In addition, the fact that the patient’s performance for the Embodied condition (and not for the Disembodied condition) were significantly different from those of the control group (Experiment 1) suggests that the effect captured in this study is unlikely to be due to an attentional bias. Fourth, the patient was aware of her deficits and was aware that we were testing her memory. We cannot exclude that the patient might have written some elements of the experiments on her diary and rehearsed them before the recall sessions. This could have helped her on some tasks (e.g. the what-where-when questions of Experiment 2, or the story content questions of Experiment 3), potentially canceling out any performance differences between conditions. However, her performances were anyway very low, and such strategy would not explain the consistent differences between Embodied and Disembodied conditions observed throughout our studies. Most importantly, it seems rather unlikely that written notes could have helped her on the perspective task. In addition, although the patient remembered some aspects of our experimental protocols (e.g. the use of VR and bodily manipulations), she did not seem to remember the details of our recall sessions.

## Conclusion

This rare case of patient, with bilateral hippocampal atrophy and related amnesia, provides critical evidence for the involvement of bodily self-consciousness in episodic memory formation. Across three experiments, we consistently show that the level of embodiment impacts the encoding of life events and the way they are later remembered. This ‘bodily self-in-memory effect’ is mediated by the hippocampus and its posterior parietal connectivity.

## Supplementary Tables

**Supplementary Table 1.** Neuropsychological tests performed in August 2022 (1 month after Experiment 1), January 2023 (1 month after Experiment 2), and December 2023 (same day as Experiment 3). RVLT = Rey Verbal Learning test, FAB = Frontal assessment battery, MOCA = Montreal Cognitive Assessment. Values written in red are out of the norms for healthy participants.

|  | August 2022 | January 2023 | December 2023 |
| --- | --- | --- | --- |
| RVLT | Delayed recall (correct):<br>9/15<br>Total recognition: 12/15<br>Total Trials 1-5: 56/75 | Delayed recall (correct): 13/15<br>Total Recognition: 15/15<br>Total Trials 1-5: 56/75 | Total Recognition: 14/15<br>Total Trials 1-5: 54/75 |
| FAB | 16 <= 16 (cut-off score) |  | 15 <= 16 (cut-off score) |
| Trail Making test | A: 33 seconds <= 81 (norm)<br>B: 79 seconds <= 299 (norm) |  |  |
| MoCA |  | 25 (norm: >= 26/30) | 26 (norm: >= 26/30) |
| Digit span |  | Raw score: 13/36<br>Standard score : 5 (norm :10) |  |
| Rey Complex Figure |  | Time to copy: 596 seconds (norm:<br>201.9s)<br>Copy: 27.5/36 (norm: 33.42) |  |
| Letter digit sequence (WAIS-IV) |  | Raw score: 4/21<br>Standard score = 3 (norm : 10) |  |
| Phonemic verbal fluency (Letter<br>P, 2 minutes) |  | Correct number of words: 17 |  |
| Semantic verbal fluency (Animal<br>category, 2 minutes) |  | Correct number of words: 8 | Correct words: 32 |
| Apathy scale |  |  | Executive Apathy: 10/24<br>Emotional Apathy: 3/24<br>Behavioral Apathy: 12/24<br>Total: 25/72 |

**Supplementary Table 2:** Experiment 2, strength of recollection. Rescaled ratings (Max:1, Min:0) of the difficulty of remembering each scene and the autonoetic consciousness questionnaire, measured one week after encoding. “Reliving E” (reliving emotions) corresponds to the question: When you think about this event now, do you re-experience any of the emotions you originally felt at the time? (100%,75%,50%,25%,0%), “Reliving G” (reliving global) corresponds to: my memory for this event is (sketchy/very detailed scale 1-7), “Visual” corresponds to: My memory for this event involves visual details (a little/a lot scale 1 to 7),”Vividness” corresponds to: When you recall this event how would you describe it in terms of vividness? (very vague/very vivid scale 1 to 7), “Location”: My memory for the location where the event takes place is (vague/very detailed, scale 1 to 7), “Spatial” corresponds to: The relative spatial arrangement of objects in my memory for the event is (vague/very detailed, scale 1 to 7), “Relived” corresponds to: Would you say you are reliving this memory or looking back on it? (reliving/looking back scale 1/0), “Rehearsal” corresponds to : How often would you estimate you have spoken about this memory since it first occurred (4-Frequently/3-Occasionally/2-Rarely/1-Never).

| Scene | Condition | Difficulty | Reliving E | Reliving G | Visual | Vividness | Location | Spatial | Relived | Rehearsal |
| --- | --- | --- | --- | --- | --- | --- | --- | --- | --- | --- |
| Printer | Embodied | 0.42 | 0.4 | 0.4 | 0.28 | 0.14 | 0.14 | 0.28 | 0 | 0 |
| Terrasse | Disembodied | 0.14 | 0.2 | 0.2 | 0.71 | 0.14 | 0.71 | 0.71 | 0 | 0 |
| Lab room | Embodied | 0.14 | 0.2 | 0.2 | 0.71 | 0.14 | 0.57 | 0.71 | 0 | 0.75 |
| Coffee machine | Disembodied | 0.85 | 0.6 | 0.8 | 1 | 0.85 | 1 | 0.85 | 0 | 0 |
| Auditorium | Embodied | 0.14 | 0.4 | 0.4 | 0.42 | 0.14 | 0.85 | 0.42 | 0 | 0 |
| Delivery dock | Disembodied | 0.14 | 0.2 | 0.2 | 0.71 | 0.14 | 0.71 | 0.71 | 0 | 0 |
| Library | Embodied | 0.85 | 0.8 | 0.8 | 0.71 | 0.85 | 0.85 | 0.85 | 0 | 0.25 |
| Reception | Disembodied | 1 | 0.8 | 1 |  | 1 | 1 | 1 | 1 | 0.75 |

**Supplementary Table 3:** Validation of BSC manipulation, Experiment 1. Scores obtained on the scoring of agency, ownership and control questions during visuomotor and perspectival match (Embodied condition) / mismatch (Disembodied condition). SD: standard deviation.

|  | Patient |  | Controls (Mean ± SD) |  |
| --- | --- | --- | --- | --- |
|  | Embodied | Disembodied | Embodied | Disembodied |
| **Agency** | 0.9 | 0.66 | 0.76±0.26 | 0.52±0.32 |
| **Ownership** | 0.91 | 0.49 | 0.76±0.26 | 0.52±0.32 |
| **Control questions** | 0.01 | 0.02 | 0.1±0.18 | 0.1±0.11 |

**Supplementary Table 4:** Experiment 1, Brain regions which functional connectivity with the hippocampus covaries as a function of the perspective score difference between Embodied and Disembodied conditions. We performed a seed-to whole brain functional connectivity using the left and right hippocampus derived from the automated anatomical labeling atlas and investigated which regions functionally connected with the hippocampus could explain the difference in performance between the Embodied and Disembodied conditions in the Perspective task (Experiment 1). The patient and the 7 control subjects were taken together for this analysis. NA : Not Applicable.

| Hippocampal seed | Directionality | Label | Cluster size | MNI coord. (x,y,z) | Cluster size pFDR | Voxel threshold |
| --- | --- | --- | --- | --- | --- | --- |
| Right | Positive | Superior Parietal Lobule left | 55 | -32 -52 +68 | 0.000046 | 0.000046 |
|  | Negative | Frontal Pole right | 51 | +12 +46 +02 | 0.000039 | 0.000039 |
|  | Negative | Angular Gyrus left | 47 | -54 -58 +46 | 0.000000 | 0.000000 |
|  | Positive | Lateral Occipital Cortex right | 35 | +46 -68 -10 | 0.000021 | 0.000021 |
|  | Positive | Superior Parietal Lobule right | 34 | +24 -50 +72 | 0.000008 | 0.000008 |
|  | Positive | Middle Frontal Gyrus left | 34 | -50 +12 +34 | 0.000009 | 0.000009 |
|  | Negative | Angular Gyrus left | 28 | -48 -54 +32 | 0.000117 | 0.000117 |
|  | Negative | Frontal Pole left | 27 | -22 +62 -04 | 0.000270 | 0.000270 |
|  | Negative | Anterior Cingulate Gyrus left | 23 | -10 +38 +10 | 0.000003 | 0.000003 |
| Left | NA | NA | NA | NA | NA | NA |

## Supplementary Figure

**Supplementary Figure 1:**
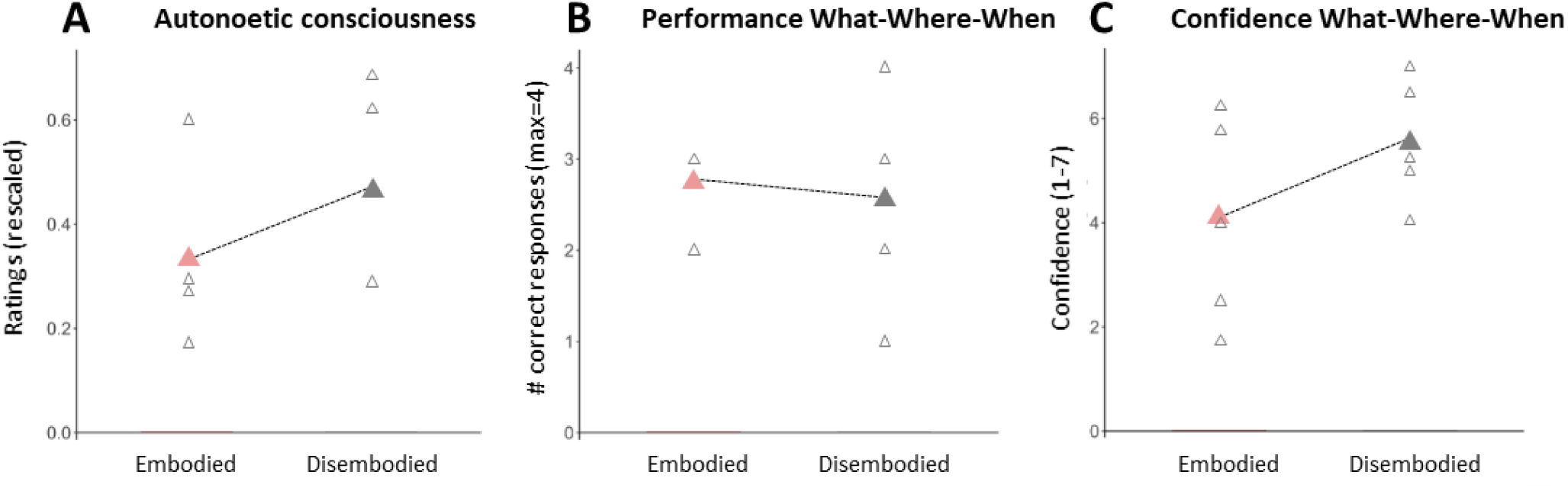
Experiment 2, Disembodied condition leads to stronger autonoetic consciousness and higher confidence. **A, Autonoetic consciousness.** The patient had a higher level of autonoetic consciousness in the Disembodied condition compared to the Embodied condition. **B, Performance What-Where-When questions.** There was no difference between Embodied and Disembodied conditions in retrieving the content and spatiotemporal context of each scene (What-Where-When). **C, Confidence on What-Where-When questions.** The patient was more confident in the “What-Where-When” answers associated with scenes encoded under the Disembodied condition compared to the Embodied condition. Empty triangles represent single trials, colored triangles indicate the mean score for each condition.

## Supplementary Text

### Methods, Experiment 3

#### Virtual Story

##### Posture I: Embodied condition, forest environment, blue guide

The patient was in the territory of a community, and had to participate in a series of rituals in order to be welcomed by the community members (**Figure 6A**). The patient was first asked to observe the forest around her (1 minute). Then, a guide (i.e. experimenter) entered the scene, wearing a blue mask, and sat in front of the patient. The patient had to reproduce the body postures that the guide would perform in front of her (ex: opening and closing the hands, with the arms extended to the front). The background voice explained the name, and meaning of the postures to the community (ex: “this posture is called “ebony trunk” and is performed as a sign of respect towards the nature”). The guide performed a series of 4 postures, which they repeated six times. Importantly, the names of the postures were invented, to ensure that the patient’s own knowledge would not interfere with the memory test.

The patient was asked to close her eyes until the next part started, to ensure a smooth transition between conditions. The guide left the scene.

##### Music I: Disembodied condition, mountain environment, orange guide

When the patient opened her eyes again she could see herself in a 3PP view, with a mountain in the background and was asked to observe the environment for 1 minute. Afterwards, the experimenter entered the virtual environment wearing an orange mask (i.e., the orange guide, see **Figure 6A**), gave a maraca to the patient, and sat with his own maraca. The patient could hear the background voice describing the characteristics of the instrument and how it was made by the community. During the first minute, the patient was invited to carefully observe it and touch it. Then, the patient had to play the instrument with the guide while music was played in the background. Four different music excerpts were played (30s each). After each music excerpt, the patient rated from 1 to 5 how pleasant that music was by shaking her instrument (ex: she shook the maraca 4 times to give a rating of 4). Then the whole series of four excerpts was repeated (no ratings). Then, the patient was asked to close her eyes, and the guide left the scene.

##### Music II: Embodied condition, forest environment, blue guide

When the patient opened her eyes again, she could see herself in the forest in 1PP. The blue guide entered the scene, gave her a sleigh bells shaker, and they performed another music ritual as before (with 4 new music excerpts).

##### Posture II: Disembodied condition, mountain environment, orange guide

The orange guide entered the scene, and they performed another body posture ritual, with a series of 4 new postures. The patient was then thanked, closed her eyes and taken back to the virtual replica of the experiment room.

The contextual story was rich in details about the rituals and the community (number and type balanced between conditions). It included many narrative elements and specific names. All of it was invented for this experiment. The forest and the mountain were two places of the same virtual environment, so they were comparable in terms of visual aspect (light, style).

Authors report no conflicts of interest

## Author contributions

- Conceptualization: NHM, MBR, JB, JP, IT, OB
- Data curation: NHM, MBR
- Formal analysis: NHM, MBR, IT
- Funding acquisition: OB
- Investigation: MBR, NHM, IT, EB, SS, BH
- Neuropsychological and neurological assessments: JP, LDPT, FE, MML, VA
- Methodology: NHM, MBR, JB, OB
- Project administration: OB, MB, NHM
- Software and virtual reality resources: FL, AT, BH, LV
- Supervision: OB
- Validation: MBR, NHM, OB
- Visualization: NHM
- Writing – original draft, NHM, MBR, OB
- Writing – review and editing: MBR, NHM, OB

*Please note that the perspective task was invented by JB during her thesis*.

## Acknowledgements

We express our gratitude to the patient for her participation in this study and for her time and patience in completing the various tasks and neuropsychological assessments, as well as to the healthy subjects who contributed to this research. We also thank Loan Mattera and Roberto Martuzzi for their expertise and assistance in acquiring the resting-state fMRI data. Finally, we extend our appreciation to Emanuela De Falco for her insightful discussions throughout the study, and to all the lab members who provided valuable advice and contributed to the technical setup of our experiments.

